# Conservation of isolate substrate preferences in mixed communities revealed through ribosomal marker protein profiling

**DOI:** 10.1101/2021.05.12.443848

**Authors:** Nicholas R Saichek, Ying Wang, Suzanne M Kosina, Benjamin P Bowen, Romy Chakraborty, Trent R Northen

## Abstract

Assessment of structure-function relationships is a central theme in microbial ecology. However, the degree that isolate metabolic activities are conserved in communities remains unclear. This is because tracking population dynamics and substrate partitioning in microbial communities remains technically challenging. Here, we describe the application of a mass spectrometry-based ribosomal marker protein profiling with stable isotope probing approach that allows for concurrent monitoring of community structure dynamics and resource assimilation within a five-member synthetic soil bacterial community. Using this approach, we find that isolate substrate preferences for glutamine and phenylalanine are largely conserved in the community and can be predicted using a weighted-sum model. However, time-series monitoring revealed a significant delay in phenylalanine incorporation by two of the strains, as well as enhanced growth for *Variovorax paradoxus* presumably due to interspecies interactions. The unique utility of this approach to temporally probe resource incorporation and community structure enables deciphering the dynamic interactions occurring within the community. Extension of this approach to other communities under various environmental perturbations is needed to reveal the generality of microbial conservation of substrate preferences.

## Introduction

Microbial communities are, in part, shaped by a dynamic pool of exogenous metabolites resulting from simple competition for substrates, symbiotic mutualisms, and complex cascades of tropic chains that are critical to sustaining earth’s ecosystems^1,2^. Yet, little is known about the conservation of microbial substrate use between isolate and community contexts because it is technically challenging to measure substrate use in mixed communities. Previously, we have observed correlations between isolates and environmental metabolites that were consistent with patterns of isolate metabolic activities, suggesting some degree of conservation of substrate use^3^. This conservation of metabolic niches would provide a powerful tool for linking community structure to overall metabolism based on the abundances and activities of constituent members^4,5^, for example to facilitate the design of synthetic communities^6^.

Prior research on microbial metabolism has mainly focused on substrate use and preferences of individual isolates grown on single or mixed substrates^7–9^. Among the limited studies exploring community resource use, mixed results have been reported regarding the relationship between metabolic function of mixed communities and that of individual community members. For example, resource use kinetics of 13 substrates (amino acids and glucose) for a three-member, co-culture were successfully predicted by a simple summation of the profiles of the three individual bacterial strains, suggesting that the presence of other species did not affect the substrate use of each individual species^10^. In contrast, analysis of carbon utilization patterns using 95 substrate Biolog plates revealed that the rate and extent of oxidation of certain, single substrates by a mixed community were not simply the sum of those exhibited by the individual isolates^11^. Deciphering the factors underlying such variability, requires assessment of the differences in metabolic behavior of individual species growing alone versus in a mixture where interspecies interactions are possible. This has been done for plant communities, with evidence showing that individual plants can change their resource use in the presence of competing neighbors with substantial ecological consequences^12^. Yet, it remains relatively unknown if such plasticity in resource use also exists among microbial communities.

Currently, several approaches exist for routine assessment of community abundance and relative community analysis by 16S rRNA amplicon sequencing is by far the most widely used despite known experimental biases. Recently, significant advancements have been implemented to overcome the extraction and amplification bias while also reducing turnaround time^13,14^.

Importantly, recent studies coupling flow cytometry with 16S rRNA sequencing as well as introduction of universal spike-in standards, generally *E. coli* 16S amplicons, prior to extraction have allowed for concurrent determination of absolute abundances of the taxa^15,16^. Metagenomic sequencing is also widely used for direct analysis of microbiome structure^17^. When combined with ^13^C-stable isotope probing (SIP)^18^, using media containing ^13^C-labeled substrates and density gradient ultracentrifugation of DNA, this approach can be used to track substrate assimilation by active microbes (DNA-SIP)^19,20^. Similarly, metaproteomics is an extremely powerful tool for measuring *in situ* protein expression within microbial communities^21–23^ and the integration with SIP enables characterization of substrate assimilation within microbial communities^24–26^.

An alternative technology that is widely used in clinical microbiology for the rapid identification of microbes is matrix-assisted laser desorption/ionization time-of-flight mass spectrometry (MALDI-TOF MS)^27–30^. Traditionally this approach involves comparison of mass spectral protein fingerprints to a database of reference spectra to generate strain identification confidence scores using a similarity coefficient-based algorithm^31^. Recently, studies have shown that changes in growth and culture conditions can alter the mass spectral patterns of the proteome, leading to overlap amongst closely related taxa^32^. To eliminate the bias attributed to this, specific efforts have focused on ribosomal proteins, as opposed to the entire proteome^33,34^. Ribosomal proteins, like ribosomal RNA (rRNA), are abundantly present and can be used as a taxa-specific diagnostic marker^35^. It has been shown that ribosomal protein profiling is suitable for analysis of simple consortia of two to three bacteria^36,37^, suggesting the possibility of using SIP in combination with MALDI ribosomal subunit protein profiling to rapidly assess both community structure and metabolic incorporation into those proteins.

Here, we hypothesized that isolate substrate preferences are conserved within a mixed community and therefore substrate use of the consortium should be accurately predicted by an isolate-informed weighted-sum model. To test this, we used a novel approach coupling of strain-specific ribosomal protein profiling with stable isotope probing by MALDI-TOF MS to monitor glutamine and phenylalanine use within a five-member bacterial synthetic community (SynCom; Fig 1). Community construction was based on our previous analysis of the translationally-active microbes present in soil samples from the Oak Ridge Field Research Center (ORFRC) in Tennessee^38^ and development of a defined medium representative of the polar small molecules found at this site^39^. Overall, using this new approach we determined that substrate use preferences were generally conserved within the five-member community, while simultaneously discovering beneficial effects attributed to interspecies interactions on the growth patterns of the community substituents.

**FIG 1.**
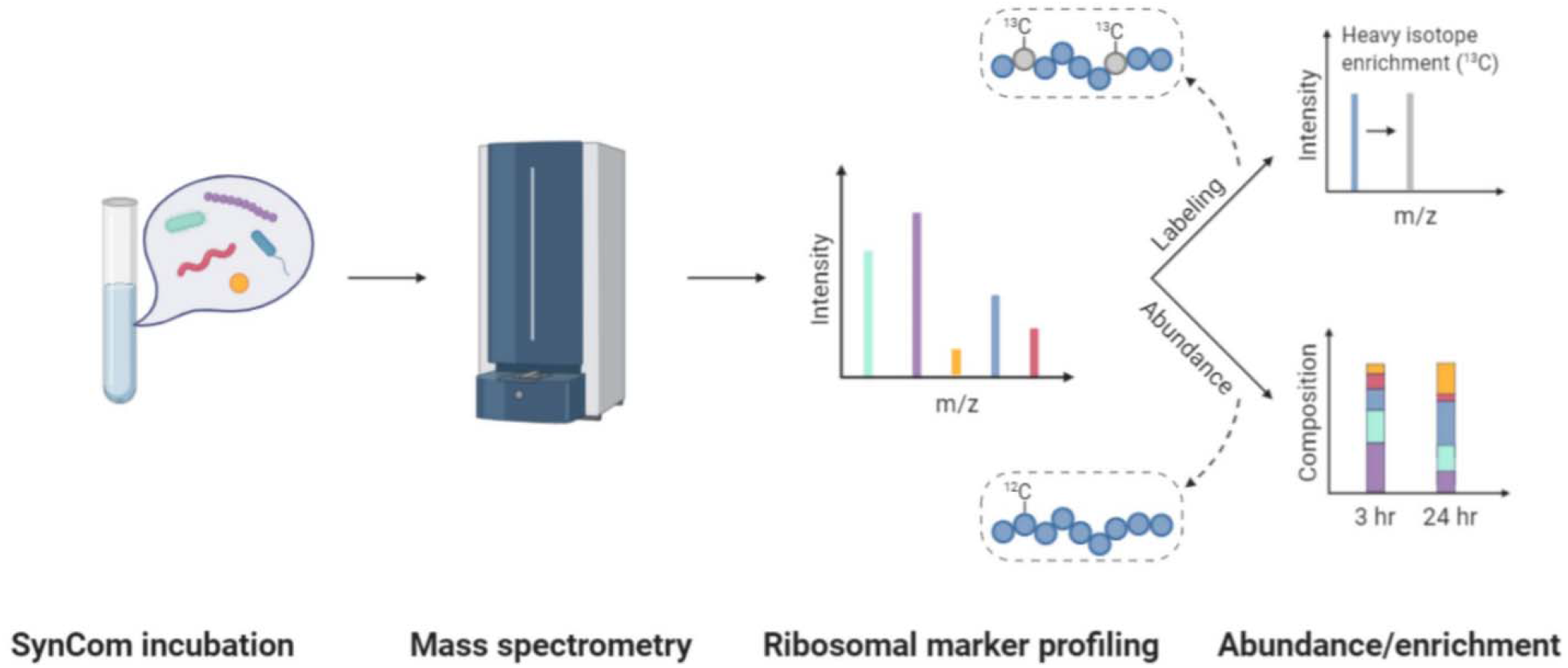
Workflow for novel mixed community SIP-proteomics approach. Schematic representation of the required steps for analysis: A synthetic bacterial community is incubated with labeled tracer metabolites. Following mass spectrometry-based ribosomal marker protein profiling, the community abundance is derived from the relative peak intensity of the respective marker proteins. Simultaneously, labeled substrate use by individual microbes within the community is measured by evaluation of ^13^C incorporation in the marker protein.

## Results

### Selection of active community isolates

The translationally-active fraction of microbial cells present in soil cores from the ORFRC were previously identified in our lab using Biorthogonal Non-Canonical Amino Acid Tagging (BONCAT) with flow cytometry and 16S rRNA sequencing^38^. Here, mapping of the 16S rRNA V4 region of isolates in our existing strain collection against the Divisive Amplicon Denoising Algorithm (DADA) sequences in the operational taxonomic unit (OTU) tables from BONCAT flow cytometry experiments revealed that, on average, 86% of the sequence reads recovered from the active community shared >97% sequence similarity with the cultured isolates from this location. From this pool of microbial isolates linked to the BONCAT flow cytometry experiments, twenty representative bacterial strains were assessed in a soil defined medium (SDM)^39^, for assembly of SynComs designed to mimic the active population. The medium was constructed in house to provide a controlled nutrient pool representative of the polar small molecules detected at the ORFRC (SI Table 1-SDM components). The concentration of sugars and amino acids was modified to reflect the ratios commonly found in soils (7:1 respectively)^40– 42^.

Sixteen of the isolates with comparable growth rates were screened for antagonistic effects during co-cultivation and plate overlay morphology analysis, further narrowing down the strain pool to eight compatible isolates. From these, five strains were selected with low average amino acid identity (AAI) similarity scores (<75%) across the proteome and then unique protein signatures (described below) were selected for downstream SynCom analyses: *Brevundimonas sp*. GW456-12-10-14-LB3, *Microbacterium hominis* FW305-3-2-15-F-LB2, *Variovorax paradoxus* GW458-12-9-14-LB2, *Pedobacter soli* GW460-11-11-14-TSB4, and *Cupriavidus necator* GW460-LB6. Plate-count titers of the SynCom cultured in SDM (inoculated at equal OD) confirmed that all strains were still present after 72 hrs of growth. Time points for analysis of early-exponential, mid-exponential, stationary, and late stationary phases of the SynCom, were identified at 3, 6, 18, and 42 hrs respectively (SI Fig 1).

### Exometabolite profiling

To determine substrate preferences of individual isolates, time-series exometabolomics (analysis of exogenous metabolite pools, also metabolic footprinting^43,44^) was performed. Specifically, monocultures grown in SDM were collected at 3, 6, 18, and 42 hr for targeted exometabolomics analysis using hydrophilic liquid interaction chromatography coupled with quadrupole time-of-flight tandem mass spectrometry (HILIC-QTOF-MS/MS). Differential substrate depletion patterns were observed among the five strains (Fig 2 and SI Fig 2). Overall, distinct groupings of depleted metabolites were observed with amino acids being generally consumed first, followed by organic acids and then nucleobases and sugars. A majority of amino acids were uniformly depleted by all strains, except *Brevundimonas sp*., within the first 6 hr of growth (Fig 2). Despite the >12-hour lag phase for *V. paradoxus* (SI Fig 1), the strain appeared to be metabolically active as indicated by the considerable consumption of a number of metabolites relative to the uninoculated control medium after 3 hr (Fig 2). Of the available carbohydrates, this strain also appeared to selectively prefer mannitol in stationary phase, whereas a decrease in monosaccharide (hexose) was observed at the later timepoints for *P. soli, C. necator*, and *Brevundimonas sp. V. paradoxus* uniquely consumed pantothenic, lactic, shikimic and vanillic acid. Unlike the other strains, *C. necator* and *P. soli* depleted a select group of nucleosides and nucleobases.

**FIG 2.**
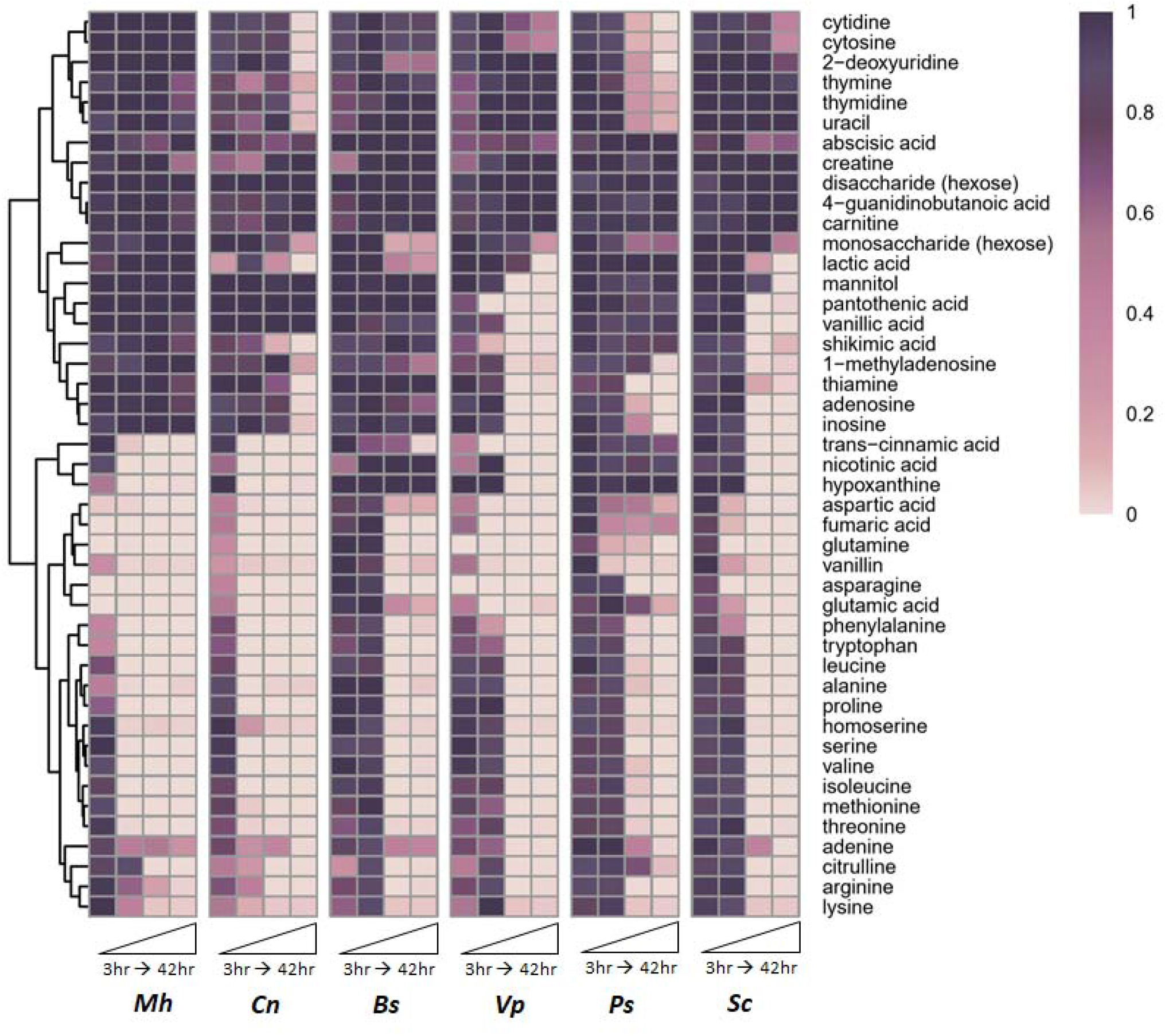
Time-series metabolite depletion patterns of monocultures and SynCom. Metabolite dynamics (52 metabolites displayed as the average peak height, normalized to the max intensity across each row) were observed in exometabolite extracts from cultures grown in a soil defined medium. Data is represented as the average of the replicates (n = 3). Strains are abbreviated as follows: *Mh-M. hominis, Cn-C. necator, Bs-Brevundimonas sp*., *Ps-P. soli, Vp-V. paradoxus*, Sc - SynCom.

### Selection and validation of strain specific ribosomal marker proteins

A SynCom-specific proteome database was constructed from spectra of each isolate across the four selected growth phases, as well as in three different defined media compositions (described in methods), to ensure robust coverage of the representative proteome. In order to assess the viability and reproducibility of strain-specific peaks as potential ribosomal marker proteins, composite summary spectra were compiled for all biological replicates. Spectral peaks present across all conditions were selected by manual matching between summary spectra. From these peaks, potential ribosomal marker proteins were identified from differential analysis between strains (*p* value < 0.05, SI Table 1 - potential marker proteins). Further, down selection of representative peak classes (list of potential ribosomal marker proteins) with an intra isolate variability in intensity <10% revealed a unique candidate marker protein for each isolate (Fig 3a).

**FIG 3.**
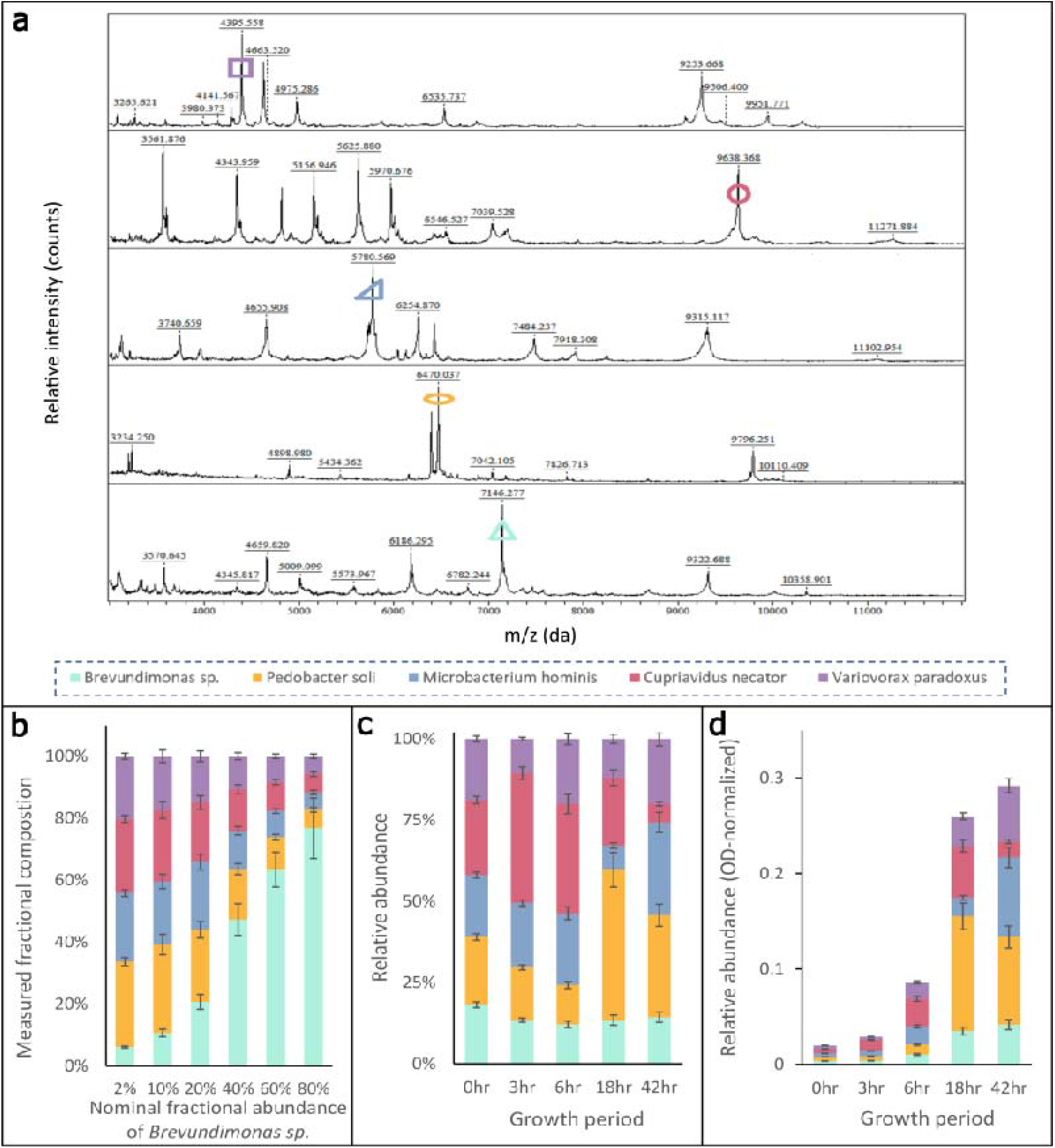
Validation and growth tracking with selected ribosomal marker proteins. **a)** Representative protein profiles for SynCom isolates; selected marker proteins are indicated by colored markers. **b)** Validation of relative community member abundance measurement using strain specific marker proteins. The fractional strain abundance for each isolate was estimated from the mean marker protein abundance in a synthetic mixture of isolates at varying concentrations of *Brevundimonas sp*. **c)** Relative abundance for each strain in the SynCom displayed over a 42 hr growth period **d)** Biomass normalized abundance during a 42 hr growth period; the y-axis represents the relative contribution of each strain with respect to the total OD of the mixed community (SI Fig 1) at each time point. Error bars are displayed as standard deviation of the mean (n = 4).

To validate the utility of these markers, encoding for specific ribosomal subunit proteins as outlined in the methods (SI Table 1-potential marker proteins), for tracking individual bacterial isolates within a community, we analyzed mixed community samples of known concentrations. Specifically, by varying the percent relative composition of *Brevundimonas sp*. from 2% to 80%, we confirmed that strain abundance could be reliably tracked with a 5% error using the selected marker proteins (Fig 3b). In order to ensure that this was true for all isolates, standard curves were performed on the individual strains. All ribosomal marker proteins showed strong correlation between their experimental and nominal percent composition as indicated by their R-squared values which were above 0.93 (SI Fig 3).

### Community composition tracking

Since the normalized peak intensities of the selected ribosomal marker proteins positively correlated with respective bacterial abundances, we used them for tracking community dynamics of the five bacterial isolates in a mixed SynCom (each strain having equal initial relative abundances and the total starting cell density of the SynCom being equal to that of the monocultures) over a 42 hr growth period in SDM (Fig 3c, 3d). To directly assess the relative abundance of each isolate in the SynCom, the community was sampled at various time points and immediately extracted for ribosomal marker protein screening. Relative abundance ratios were validated using triplicate plate titer counts of serial dilutions 10^4^ to 10^6^ cfu/mL (SI Table 1-titer counts). As can be seen from the community composition plot (Fig 3c), *C. necator* was dominant in the community early on (before 6 hr), whereas *P. soli, M. hominis*, and *V. paradoxus* became dominant by 42 hr. While *V. paradoxus* did not show any growth until 12 hr in monoculture (SI Fig 1), considerable growth was observed in the SynCom after just 6 hr which suggests beneficial interactions within the community.

### Modeling resource use in monocultures and community

Since community structure suggested possible interspecies interactions, we wanted to test the conservation of substrate use in a mixed community compared with that in monoculture. To do this, we specifically focused on two amino acids, glutamine and phenylalanine; both are routinely detected in soil water and can serve as carbon and nitrogen sources for soil microbes^3,39,45^. These two substrates showed varying depletion patterns in the exometabolite profiles by the five monocultures (Fig 2 and SI Fig 2). The time-resolved metabolite depletion data (Fig 2) were fitted to a previously published model by Behrends et al.^46^. Specifically, the depletion kinetics of individual substrate from a mixture of substrates by an isolate over the course of growth in batch culture condition can be described by a sigmoid function (Methods, Equation 1). We found for each of the five bacterial monocultures, their depletion of glutamine and phenylalanine followed the Behrends’ model with an average R-squared value of 0.998 (SI Fig 4, 5 and Table 1-estimated modeling parameters). Then the measured depletion data and fitted depletion kinetics were subtracted from the initial concentrations of the substrates in the medium and converted to usage data and simulated usage kinetics for each monoculture (Fig 4a). The two parameters describing the substrate usage properties—T_50_ (the time when half of the available substrate has been used) and width (the duration when the substrate is used at the maximum rate)—differed significantly among the five bacteria (Fig 4b, SI Fig 6), demonstrating differential substrate use preferences. *M. hominis* depleted phenylalanine fastest, with the smallest T_50_ value (2.81 ± 0.07 h), followed by *C. necator* (3.39 ± 0.14 h), *V. paradoxus* (4.35 ± 0.13 h), and finally *P. soli* (8.22 ± 0.27 h) and *Brevundimonas sp*. (8.61 ± 0.30 h; Fig 4b, right). As for glutamine, *M. hominis* (2.17 ± 0.16 h) and *V. paradoxus* (2.10 ± 0.04 h) depleted glutamine faster than *C. necator* (2.79 ± 0.01 h) and *P. soli* (2.94 ± 0.03 h), with *Brevundimonas sp*. being the slowest (10.86 ± 0.37 h; Fig 4b, left). As the T_50_ values increased, the time window around maximum uptake rate (width) also extended (SI Fig 6).

**FIG 4.**
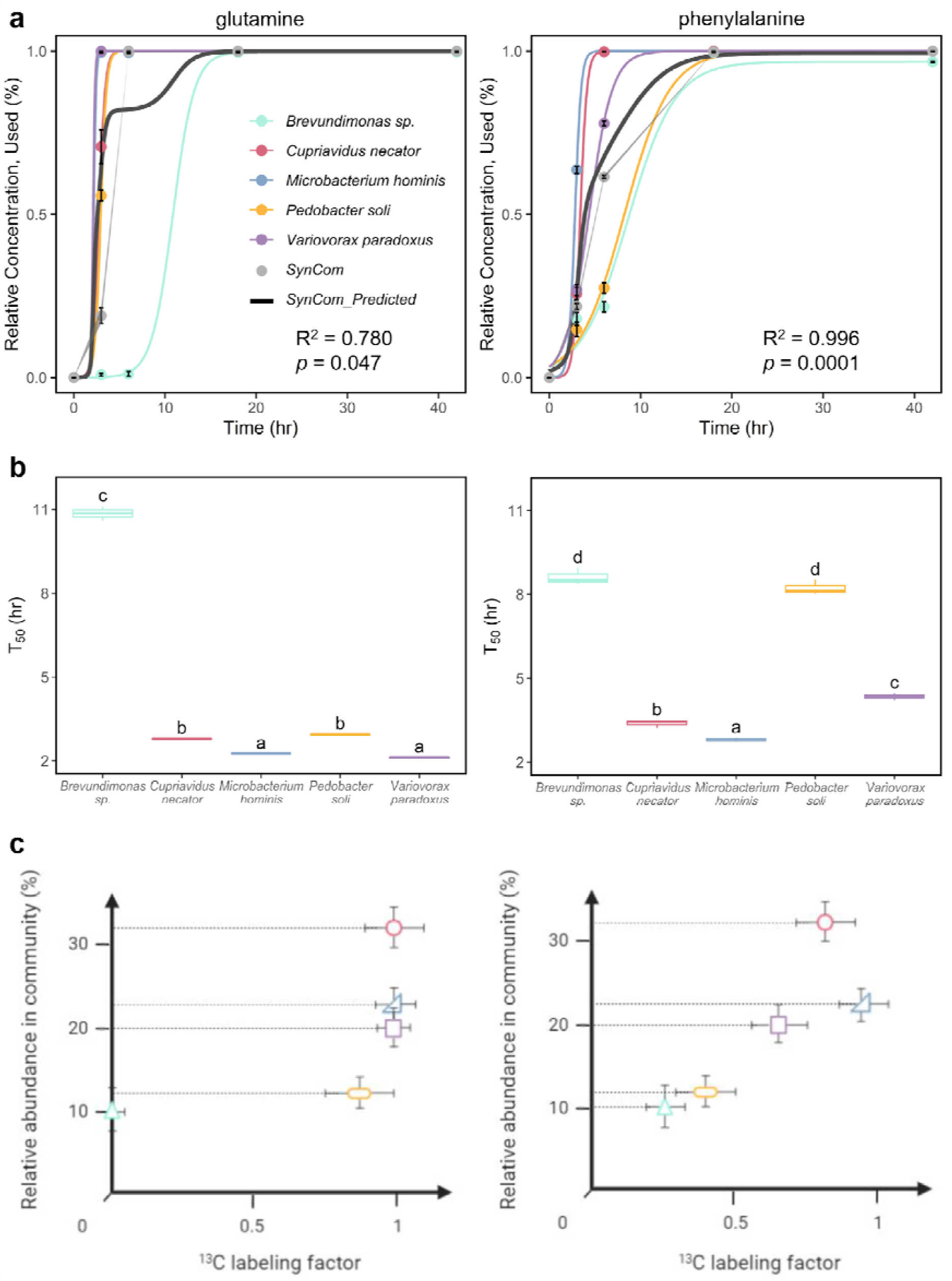
Modeling vs measured substrate usage in monocultures and SynCom. **a)** For each bacterial species grown in monoculture, their time-resolved usage data (colored markers) of glutamine (left) and phenylalanine (right) were fitted by a previously published model by Behrends et al. (2009) using nonlinear least squares (SI Fig 4, 5). The five simulated monoculture usage kinetics (colored lines) were summed and weighted by the starting relative abundance of each species in the SynCom to produce the predicted total usage curve of the five-member SynCom (SynCom_Predicted: dark gray line). The light gray circles represent the measured substrate usage by the SynCom. Error bars (for five monocultures and SynCom) and shading (for SynCom) represent standard error of mean (n = 3). **b)** Estimated T_50_ values for the use of glutamine (left) and phenylalanine (right) for each monoculture. Letters signify significant differences (*p* < 0.05) between species for the same substrate, using one-way ANOVA with post-hoc Tukey HSD test (n = 3, except for glutamine n = 2 for *Brevundimonas sp*., *C. necator*, and *P. soli*; SI Fig 4, 5, Table 1). **c)** Incorporation of ^13^C-glutamine (left) and ^13^C-phenylalanine (right) labeled substrates into strain-specific ribosomal marker proteins in the mixed community after 6 hr growth. The relative abundance in the community is derived from the y-axis in Fig 3c at 6 hr, whereas the ^13^C labeling factor is a measure of the relative mass shift. Error bars are represented as standard deviation of the mean (n = 3).

**FIG 5.**
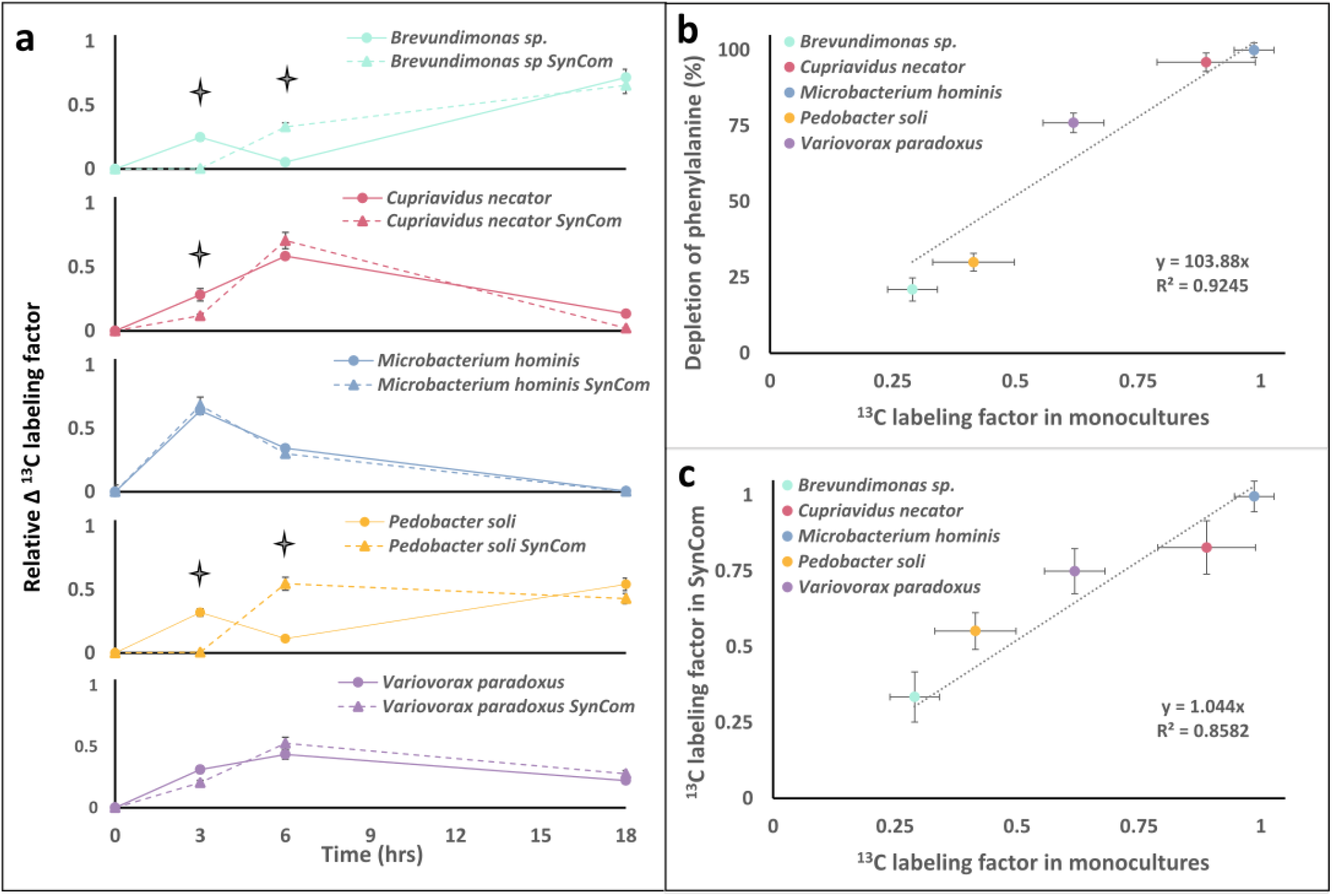
Comparative usage of ^13^C-phenylalanine. **a)** Incremental usage of ^13^C-phenylalanine by strains in monocultures and SynCom determined by ribosomal marker protein SIP. Significant differences, from one-way ANOVA, in usage between monoculture and SynCom are denoted by ✦ (*p* < 0.05), where error bars represent standard error (n = 3). **b)** Plot for phenylalanine use by monocultures at 6 hr as determined by exometabolomics (Fig 2) and cumulative ^13^C-enrichment in ribosomal marker protein. **c)** Comparative plot of overall usage of ^13^C-phenylalanine by individual strains in monocultures and SynCom by 6 hr. Error bars are represented as standard deviation of the mean (n = 3).

To predict the usage kinetics of the five-member SynCom, the monoculture usage curves were summed and weighted for the starting relative abundance of each species in the community (Fig 3c). The modeling predictions were then compared with measured substrate usage dynamics by the SynCom determined by exometabolomics profiling (Fig 2). We found different trends for the two substrates (Fig 4a). The predicted SynCom usage of phenylalanine agreed well with the measurement (R^2^ > 0.99, *p* = 0.0001; Fig 4a, right). However, glutamine usage measured in the SynCom deviated from model prediction (R^2^ = 0.78, *p* = 0.047; Fig 4a, left) and showed near complete depletion by 6 hr, whereas the model predicted only 82% of use at this timepoint.

### Tracking resource incorporation into ribosomal marker proteins

To further evaluate how the substrate was partitioned amongst its members, we used stable isotope probing in combination with ribosomal marker protein profiling. Previous studies have focused on endpoint analysis to probe metabolic flux by measuring the change in isotope incorporation^47^. Here, we chose three isotopically-labeled substrates, ^13^C-glutamine, ^13^C-phenylalanine, and ^13^C-glucose, to track relative substrate incorporation by each member within the SynCom over time. To do this, the five bacterial strains were grown in SDM, with the ^12^C substrate replaced by the ^13^C-labeled isotopologue, as both monoculture and a five-member mixed community. Incorporation of these labeled substrates into the ribosomal marker protein was estimated by annotating the average mass shift of the protein as a result of heavy isotope enrichment^48^.

The usage for subsequent enrichment studies was determined by comparing the observed mass to the theoretical mass for each respective marker protein. We estimated the upper labeling bound by growing monocultures in ^13^C-glucose, a preferred bacterial carbon source (SI Fig 7). For glutamine and phenylalanine, differential ^13^C incorporation in ribosomal marker proteins was observed by different strains in the SynCom at 6 hr (Fig 4c). Specifically, in ^13^C-glutamine-labeled medium, *Brevundimonas sp*. showed no observable mass shift (in the strain-identifying marker protein) but markers for the other four SynCom members all showed considerable enrichment; this was expected given the delayed glutamine utilization by *Brevundimonas sp*. in monoculture (Fig 2 and SI Fig 2).

Interestingly, analysis of the incremental usage of ^13^C-phenylalanine over each time range revealed dramatic temporal shifts in usage patterns between the SynCom and monocultures (Fig 5a). We observed that in both monoculture and consortia *M. hominis* rapidly incorporated phenylalanine in the first 3 hr, whereas other strains showed delayed consumption of the substrate in the SynCom. Specifically, *P. soli* and *Brevundimonas sp*. incorporated little or no ^13^C in the SynCom with respect to their incorporation levels in monocultures during the first 3 hr. However, they both showed increased incorporation during the 3-6 hr period. There was also a considerable difference for *C. necator* during the 6-18 hr period. In monoculture, this strain appeared to incorporate a greater amount of the phenylalanine during the 6-18 hr period than it did in the SynCom, suggesting that consumption in the SynCom had diverted to other substrates.

## Discussion

Resource utilization by microbial species with varying nutritional needs is important in structuring microbial communities and ultimately determines overall community metabolic activity^49,50^. In this study, we examined the conservation of microbial substrate preference for glutamine and phenylalanine in monoculture vs SynCom. This was done using strain-specific ribosomal marker protein profiling combined with stable isotope probing to simultaneously measure relative strain abundance and substrate incorporation into a five-member bacterial SynCom.

For monocultures, differential substrate preferences were observed (Fig 2 and Fig 4). Specifically, for all five isolates except *Brevundimonas sp*., glutamine was preferred over phenylalanine, as indicated by the smaller T_50_ values reflecting earlier depletion during the growth. Similar trends have been reported previously for other bacteria, specifically two Pseudomonads and one Bacilli isolated from groundwater^10^. A marine bacterium, *Pseudoalteromonas haloplanktis*, was also found to use glutamine as a preferred amino acid source, with >95% depletion after 4.5 hr of growth, while phenylalanine depletion was slow during the first 3 hr of growth before being depleted to negligible concentration by ∼7.5 hr^51^. Genomic analysis suggests that the delayed use of glutamine by *Brevundimonas sp*. vs other strains may be due to the lack of the gene encoding for glutaminase (EC 3.5.1.2)^52^.

To evaluate if isolate substrate preferences are conserved in the mixed community, we applied a weighted-sum model to predict community substrate use. Ecological theory suggests that if no interspecies interactions are present, the functioning of a mixed community should be equal to the combined output of the constituent components; deviations from this would imply interspecies interactions^53,54^. The overall resource use by the five-member SynCom was predicted from the usage kinetics of each individual isolate using a summation model weighted by the starting relative abundance of each member^54^.

For phenylalanine, the usage kinetics by the SynCom was accurately fit using the weighted-sum resource model, suggesting that interspecies interaction did not affect the use of phenylalanine by each species in the SynCom. Analysis of the ^13^C-phenylalanine incorporation into ribosomal marker proteins for each member within the SynCom also supported this inference. Specifically, strong correlation was observed between phenylalanine use, quantified by isolate exometabolomics, and ^13^C labeling factor for each strain-specific marker protein after 6 hr growth in ^13^C-phenylalanine-labeled medium (Fig 5b). Moreover, the degree of phenylalanine incorporation by the five bacteria in the SynCom was proportional to that in monocultures for the first 6 hr (Fig 5c), indicating the behavior of each strain with respect to phenylalanine was mostly conserved in the mixed community. This is interesting given that there are likely many confounding processes resulting from substrate co-metabolism and distribution throughout the proteome occurring simultaneously. One plausible reason ribosomal marker protein-SIP served as an accurate proxy for substrate use, at these early time points, could be in part due to the rapid turnover of ribosomal proteins in prokaryotes^55^.

For glutamine, the predicted usage was slower than the actual overall community usage measured by exometabolite profiling. This deviation could be due to interspecies competition for this preferred amino acid source (leading to rapid decrease in concentration) or use of alternative substrates produced by community members^56,57^. Future analysis of the ^13^C-spent media could provide useful insight in examining the rates of metabolite depletion in the SynCom. In monoculture, four of the strains depleted almost all the glutamine (>99%) within the first 6 hr with an average T_50_ of 2.5 hr (Fig 2, 4a, 4b). In particular, with the smallest T_50_ among the five species, *V. paradoxus* depleted glutamine within 3 hr, long before there was any appreciable growth (SI Fig 1). This suggests that this strain tended to preferentially accumulate glutamine without replication. Similar findings have previously been reported; for example, *Bacillus cereus* depleted glucose during early growth stage, likely due to a significant delay in substrate conversion during replication or rapid transformation of glucose into other compounds such as glycogen^10^. When examining the community structure measured using unique ribosomal marker proteins over time, we observed more substantial growth for *V. paradoxus* in the SynCom than in monoculture at 6 hr (Fig 3c, 3d), indicating its promoted competitive advantage due to interspecies interactions^50^. Taken together, the preferential accumulation of glutamine by *V. paradoxus* and its increased growth in the SynCom could have led to the faster usage of glutamine in the community than the model prediction.

These findings have implications for how resource utilization and competition can structure bacterial communities. Niche theory states that one mechanism for species to stably coexist is to increase niche differentiation via different patterns of resource use^58,59^. Our previous work reported little overlap in exometabolite niches among sympatric microbes in two soil environments^8^. In this work, likely because our soil defined medium contains a relatively small number of common bacterial substrates, more overlap in substrate use was observed between the five species. When competing for overlapping resources, we found isolate substrate use preference was mostly conserved and determined the relative partitioning of use within the community. This agrees with contemporary resource competition theory which states that monoculture resource utilization traits can predict resource competition in mixed cultures^60^. Yet this did not preclude the presence of metabolic interactions among species and impacts on the community. For phenylalanine, the R-squared fit of our weighted-sum model to predict community usage was > 0.99, but we still observed differences in temporal substrate usage as compared with monoculture. For glutamine, with a higher level of usage overlap among majority of the community members, the overall use was not just an aggregated term, but also dependent on the competitive abilities of the constituent species within the community setting^61^, as mentioned above for *V. paradoxus*. We speculate that dynamic metabolic adaptation to maximize the relative fitness under competition^62^ and/or facilitative interactions, such as co-metabolism of waste products by other species^56^, may have improved the competitive advantage of certain species within the community. Future work combining our ribosomal marker protein-SIP approach with integrated modeling of community structure and substrate use will enable a more mechanistic understanding of such interactions.

Overall, we find that the adaption of protein profiling, which is widely used in clinical applications^30,63^, and to our knowledge has not been used for environmental studies, enables rapid profiling of community structure and resource incorporation partitioning within a SynCom. This enhanced capability for performing parallel analysis of ^12^C-media, for community abundance profiling, and ^13^C-media, for resource partitioning, affords additional benefits in comparison to the conventionally utilized 16S rRNA amplicon sequencing. However, to facilitate broader application of this approach for the research of resource-based interactions in microbial communities, a few considerations need to be addressed. First, selecting unique ribosomal marker proteins for each microbe in a community requires prescreening or construction of a ribosomal marker protein reference database and it is unlikely that this approach can be effectively applied to communities consisting of more than approximately 30 strains. Second, the inherent difficulty in accounting for primary vs secondary users, as well as incorporation of the label into untracked parts of the proteome, adds additional complexity to the perceived degree of labeling occurring. To limit the effects from this, incubation times were kept relatively short, but this adversely resulted in some peak broadening due to the vast degree of labeling occurring simultaneously. Although using intact proteins, as opposed to peptides, did convolute the isotopic envelope distribution measurements^26^, the rapid and robust nature of this approach was sufficient for determination of the role resource competition and cross-feeding play in the structuring of microbial communities. However, we see significant opportunities to integrate this approach with full metaproteomics^23,24^ to provide a more complete understanding of community activities. Specifically, the rapid throughput of ribosomal protein profiling lends itself to exploring large experimental spaces (e.g., substrates, environmental conditions, etc.) which can be examined in great detail using slower but more comprehensive metaproteomic methods.

In conclusion, using stable isotope probing of strain specific ribosomal proteins we found that isolate substrate preferences for glutamine and alanine were largely conserved in mixed communities and were able to predict substrate use for a mixed community of five bacteria from isolate cultures. Together this integrated modeling and testing framework provides a platform for designing and refining synthetic communities based on knowledge of substrate preferences and analysis of ribosomal protein markers. We anticipate that this will provide an important new tool for gaining insights into metabolite uptake and exchange within laboratory consortia and if deployed broadly across diverse organisms could define the extent of microbial substrate conservation in isolation vs consortia.

## Materials and Methods

### Chemicals

Glucose (CAS 50-99-7) was from Amresco (Solon, OH). ^13^C-glutamine (C5-99%), ^13^C-L-phenylalanine (ring, C6-99%), ^13^C-glucose (C6-99%), and algal amino acid mixtures were from Cambridge Isotopes Laboratory, Inc. (Tewksbury, MA). LCMS-grade methanol and water were from J.T. Baker (Avantor Performance Materials, Center Valley, PA). All other standards, amino acid kits, and metabolites were from Sigma-Aldrich (St. Louis, MO).

### Bacterial isolates

A bacterial isolate collection of strains isolated from the Oak Ridge Field Research Center at U.S. Department of Energy Oak Ridge National Laboratory (Oak Ridge, TN) was provided by Romy Chakraborty and Adam Deutschbauer (Lawrence Berkeley National Laboratory, Berkeley, CA). Isolates were revived from glycerol stocks on SDM medium for use in culture experiments. Lab isolates with similar growth rates were selected for SynCom model construction based on closest 16S match to the most abundant sequences identified in the active fraction of Oak Ridge soils using BONCAT labeling^38^.

### Media Construction

Oak Ridge Field Research Center soil defined medium (SDM) was constructed as previously described with slight modifications^39^. The base media contained the same 46 metabolites observed in the metabolite quantification experiments of soil from the ORFRC field site (SI Table 1-SDM components), with the concentrations altered to reflect the sugar: amino acid ratios commonly observed in environmental samples^40–42^. An additional metabolite medium containing 14 essential organic acids each at 5 µM along with Wolfe’s vitamin and Wolfe’s mineral solutions at 1X concentration were added to the base media^64^. Growth of 20 BONCAT active representative field isolates was screened in SDM at 30 °C for 42 hr to select candidate strains with similar growth rates for construction of the five-member SynCom. Agar plates for solid culture assays were prepared with 1.5% agar.

### Co-culture phenotype screening

Bacterial overlay plating with microarray stamping was performed for compatibility screening of the isolates^65^. Briefly, overnight cultures were spread as a thin layer on SDM agar plates and allowed to air dry before two microliters of pure isolates were individually deposited in an overlay technique. The cultures were spotted 10 mm apart in a 6 x 6 microarray on a square petri dish to ensure the cultures were spatially separate. Clearing zones and growth morphologies were observed after a 42 hr incubation period at 30°C to determine interaction trends.

### Isolate growth and extraction studies

Monocultures of the SynCom strains were revived from freezer stocks (30% glycerol) into SDM at 30 °C for 18 hr. After incubation, cultures were centrifuged at 3800 x g for 5 min and washed with PBS (1X, pH 7.2). Cells were resuspended in fresh SDM and the OD_600_ was adjusted to 0.05 ± 0.025 for initialization of growth studies. Readings at OD_600_ were taken every 15 minutes with orbital mixing at 30 °C for 72 hr for precise determination of cell density and lag phase using a Biotek Synergy H1m monochromator-based multi-mode microplate reader. Growth was defined as an increase in OD of greater than 0.05 from the initial time point, after media control subtraction. Aliquots were removed at 3, 6, 18, and 42 hr for downstream exometabolomic and ribosomal marker protein processing. For exometabolite profiling, 200 µL of bacterial culture was centrifuged at 3800 x g for 5 min in triplicate, followed by filtration of the supernatant through a 0.22 μm PVDF spin filter (MilliporeSigma, Burlington, MA) to remove cell debris, and then lyophilized. Dried samples were resuspended in 150 μL of LCMS-grade methanol, spiked with a labeled internal standard mix consisting of amino acids, nucleosides, nucleobases, and sugars spanning the reported time window, and stored for LCMS analysis (as described below). Parallel culture aliquots were normalized to a final OD_600_ of 0.5 before quenching for ribosomal protein profiling. For quenching and subsequent MALDI-TOF MS analysis, cells were harvested by refrigerated centrifugation (10 °C) of OD-normalized cultures at 3800 x g for 5 min followed by washing with PBS. Cell pellets were resuspended in 50 μL of a 70% ethanol/1% TFA solution with 30 mg/mL thymol, added prior to incubation at 52 °C for 10 min, in order to simultaneously quench the metabolism and extract the proteins. After heating, the permeabilized cells were stored at -20 °C for MALDI-TOF MS analysis (as described below). Nutrient agar plates were used for serial dilution colony titer assays to validate relative species abundance. All strains displayed unique colony morphologies, under the outlined growth conditions, allowing for accurate plate counts.

### Ribosomal protein marker selection and validation

A database was constructed using composite summary spectra for each of the strains in the SynCom for ribosomal marker protein selection. Briefly, each isolate was grown in three minimal media (SDM, Wolfe’s Modified Minimal media, and 14AA basal medium; SI Table 1) and aliquots were removed at various time points for downstream processing to ensure the selected ribosomal marker proteins were not phase or metabolite composition dependent; these media and time conditions are reflected as biological replicates for data processing comparisons. Raw spectra (n = 36; per isolate) were imported and processed, as described below in data analysis, to select peaks with a signal to noise ratio greater than 10. The technical and biological replicates were compiled into composite spectra using a similarity filter of 95 and 75% respectively. Multidimensional scaling (MDS) was used to visualize the spectral similarity and ensure distinct groupings of the isolates were present. Two-way clustering was performed on the data matrix which allowed for potential strain-level ribosomal marker proteins to be defined; peak classes were selected with a p value < 0.05 (SI Table 1-potential ribosomal marker proteins). The defined ribosomal marker proteins were screened against twenty unknown spectra to ensure correct strain identification was achieved (>95% similarity score). To determine the identities for the protein markers the genome assemblies were subjected to *in silico* extraction of the nucleotide sequences using tBLASTn in KBASE prior to monoisotopic molecular weight prediction using the ExPASy protein parameter tool^52,66^. The most common posttranslational modifications, specifically N-terminal methionine loss, phosphorylation, and methylation were taken into consideration to ensure monoisotopic mass predictions were accurate.

### Community abundance and resource utilization

Mixed communities were inoculated in a similar fashion as the growth studies. Briefly, isolates were revived from freezer stocks and washed after overnight incubation in SDM. For abundance and utilization studies, isolates were mixed equally (OD normalized) into fresh media and incubated for 3, 6, 18, and 42 hr prior to metabolic quenching. Protein samples were spiked with ubiquitin (Sigma-Aldrich BioUltra, 5 nmol), a small subunit protein native to eukaryotes, was added prior to quenching as an internal standard to account for instrumental variance between samples. Relative abundance measurements were performed by comparing the relative ratios of the marker protein peaks in the mixed community at various time points. Isotope enrichment studies were performed using identical media components, with the substitution of a single ^13^C-labeled substrate for its ^12^C-counterpart. Three metabolites of interest, ^13^C-glutamine (C5-99%), ^13^C-L-phenylalanine (ring, C6-99%), ^13^C-glucose (C6-99%), were screened over the duration of the growth phase for each individual isolate to ensure incorporation into the ribosomal marker protein was feasible.

### LCMS analysis

Exometabolite extracts were chromatographically separated by hydrophilic liquid interaction chromatography (HILIC) on an Agilent 1290 series HPLC system equipped with an InfinityLab Poroshell 120 HILIC-Z, 2.1 × 50 mm, 2.7 µm, PEEK-lined column (Agilent 679775-924) employing a 7 min gradient. Liquid flow was directed to an Agilent 6550 quadrupole time-of-flight tandem mass spectrometer (Q-TOF MS/MS) operated in positive and negative polarity for analysis. All parameters and timetables are outlined in SI Table 1-LCMS operating conditions.

### MALDI-TOF MS analysis

Each aliquot was mixed with sinapinic acid (10 mg/mL in 30% acetonitrile/1% TFA) in a 5:1 ratio (matrix:sample) prior to spotting one microliter in triplicate spots on the ITO-coated glass slide. The final solution was spotted in triplicate as one microliter spots. Technical replicate spectra were independently acquired using a Bruker Ultraflextreme MALDI-TOF MS (Bruker Daltonics; Billerica, MA) equipped with a 337 nm Nd:YAG laser. Spectra were manually acquired as 500 shot composites in positive linear mode within a mass range of 3 to 25kDa using FlexControl software (version 4.0; Bruker Daltonics; Billerica, MA). Instrument calibration was performed using the Bruker Protein Calibration Standard I mixture according to manufacturer specifications.

### Data analysis and modeling

Metabolomics files were converted to mzML format (ProteoWizard MSConvert, Palo Alto, CA)^67^ and analyzed using Metabolite Atlas (https://github.com/biorack/metatlas) in conjunction with Python to construct extracted ion chromatograms (EIC) corresponding to metabolites contained in SDM and our in-house standards library^68,69^. Authentic standards were further used to validate assignments based on accurate mass less than 15 ppm, retention time within 0.1 min of predicted, as well as MS/MS matching of major fragment ions (SI Table 1-metabolite identifications). Internal standards and quality control mixtures were assessed to ensure no sample-to-sample retention time drift or reduction in intensity occurred; all metabolites that eluted before the solvent front (*t*_*o*_ = 0.2 min) were excluded from the analysis. Peak height tables were processed and used to generate heatmaps for direct metabolite usage comparison. Statistical analyses were performed using Microsoft Excel and R v3.6.1 based on a p-score <0.05. Raw data and processed outputs have been deposited at the Joint Genome Institute’s Genome Portal under Project ID: 1280078 (https://genome.jgi.doe.gov/portal/202ics_FD/202ics_FD.info.html).

To model glutamine and phenylalanine depletion by each individual species grown in monoculture, substrate relative abundance data (*s*) measured by metabolomics over time (*t*) were fitted to the Behrends’ model^46^, using nonlinear least squares in R v3.6.1:

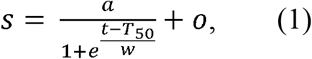

where *a* is the amplitude, calculated as the difference between the starting and the lowest level of substrate abundances; *o* is the offset, defined as the lowest substrate abundance; *T*_50_ is the half-life, defining the time at half-amplitude; and *w* is width, defining the time window when the exponent of *e* to go from 1 to -1, roughly showing the “uptake window” when the substrate is depleted at the maximum rate.

Raw mass spectra were exported as ASCII files using FlexAnalysis software (version 4.0; Bruker Daltonics; Billerica, MA) and imported into BioNumerics software (version 7.6, Applied Maths, Sint-Martens-Latem, Belgium) for ribosomal marker protein selection and evaluation. Raw mass spectra were smoothed with a Kaiser Window of 20 pts, baseline subtraction was performed with a rolling disc algorithm size of 75 pts, noise was computed using a continuous wavelet transformation (CWT) with a s/n cutoff of 10. Hierarchical classification was performed using Spearman ranks and Euclidean distances. For ^13^C-enrichment studies, heavy isotope enrichment was measured by comparing the observed mass increase to the predicted nominal mass increase of the uniformly labeled protein. Temporal usage was calculated as follows to determine usage of labeled substrate with respect to non-labeled media components:

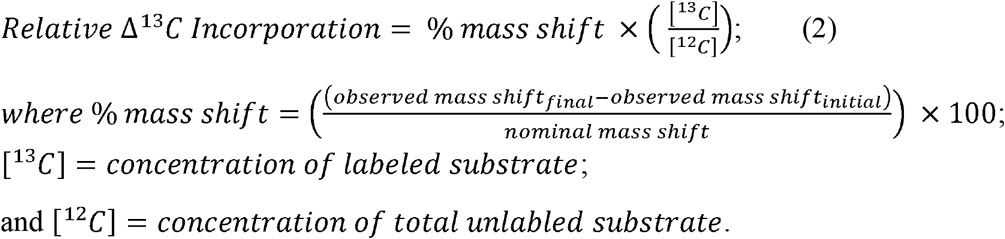

## Author contributions

*NRS and TRN conceived the study and designed the experiments. NRS and YW wrote the manuscript. NRS performed the experiments. NRS and SMK analyzed the metabolomics data. NRS and YW designed the figures. BPB and YW developed code for modeling and data interpretation. RC isolated and sequenced all strains used in study. All co-authors commented on the design of experiments, data analysis and draft manuscripts*.

## Acknowledgments

*This material by ENIGMA-Ecosystems and Networks Integrated with Genes and Molecular Assemblies (http://enigma.lbl.gov), a Science Focus Area Program at Lawrence Berkeley National Laboratory is based upon work supported by the U*.*S. Department of Energy, Office of Science, Office of Biological & Environmental Research under contract number DE-AC02-05CH11231*.

## Supplementary Information

SI Table 1: Supplementary Excel file (media, metabolites, strains, marker proteins, modeling)

**SI Fig 1.**
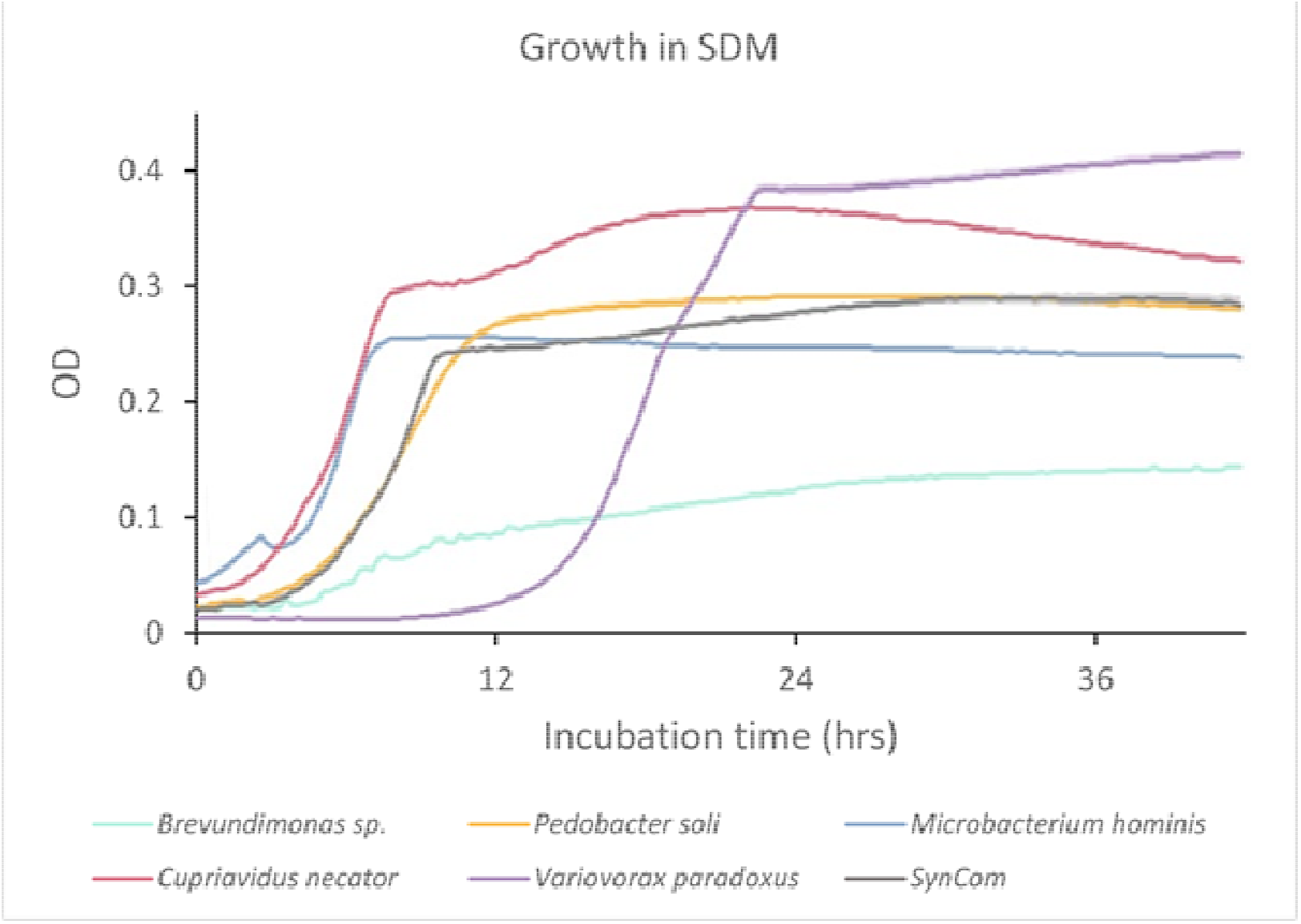
Growth curves for monocultures and SynCom in soil defined medium (SDM) over 42 hr. Optical density readings were taken on a Biotek Synergy H1m monochromator-based multi-mode microplate reader, λ = 600nm and a path length of 4.2mm. Relative standard deviation (RSD; n = 3) for all strains was <5% and is represented by the shaded area.

**SI Fig 2.**
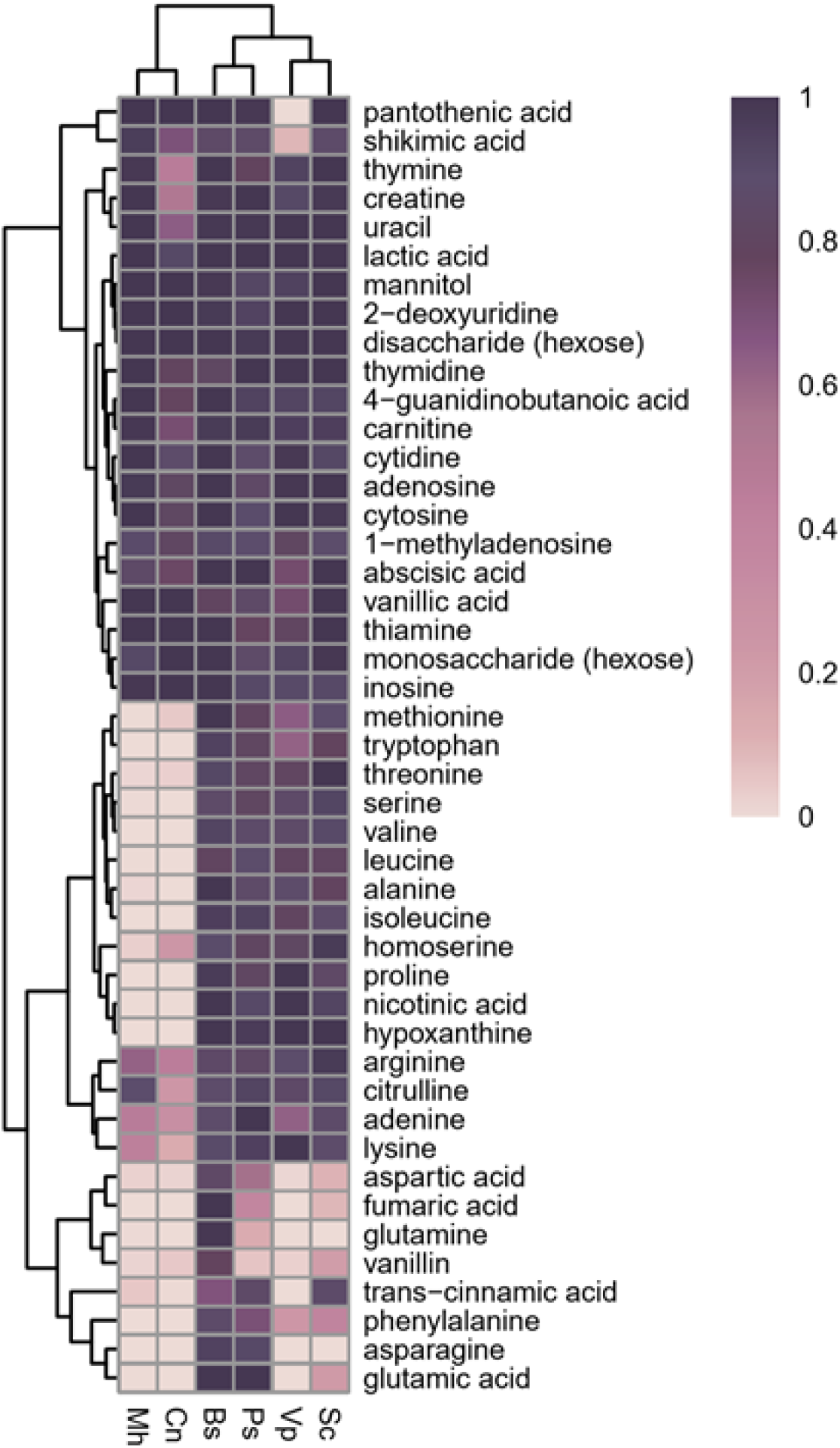
Exometabolite depletion by monocultures and SynCom in SDM after 6 h. Relative metabolite abundances were measured using LC-MS and max intensity of each metabolite was normalized relative to the media control. Strain designations are abbreviated as follows: *Mh-M. hominis, Cn-C. necator, Bs-Brevundimonas sp*., *Ps-P. soli, Vp-V. paradoxus*, and *Sc-*SynCom. Data is represented as the average of the replicates (n = 3).

**SI Fig 3.**
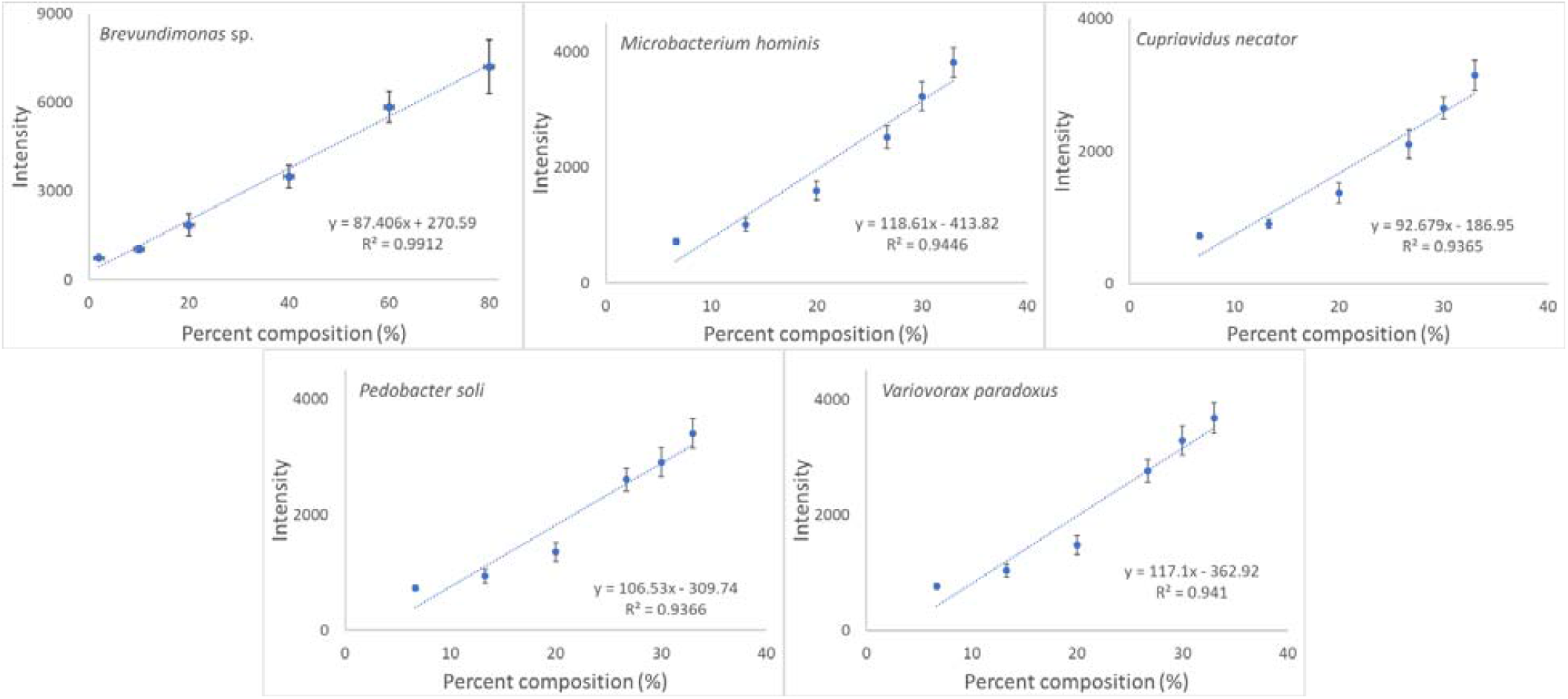
Validation of ribosomal marker proteins in a five-member mixed community. Isolates were grown overnight as monocultures in SDM. Cultures were washed and normalized to 0.1 OD (600 nm) before dilution and mixing to defined community compositions. Individual proteins were used as markers for each strain and tracked using MALDI-TOF MS to determine the relative ratios. Error bars represent standard deviation of the mean (n = 4).

**SI Fig 4.**
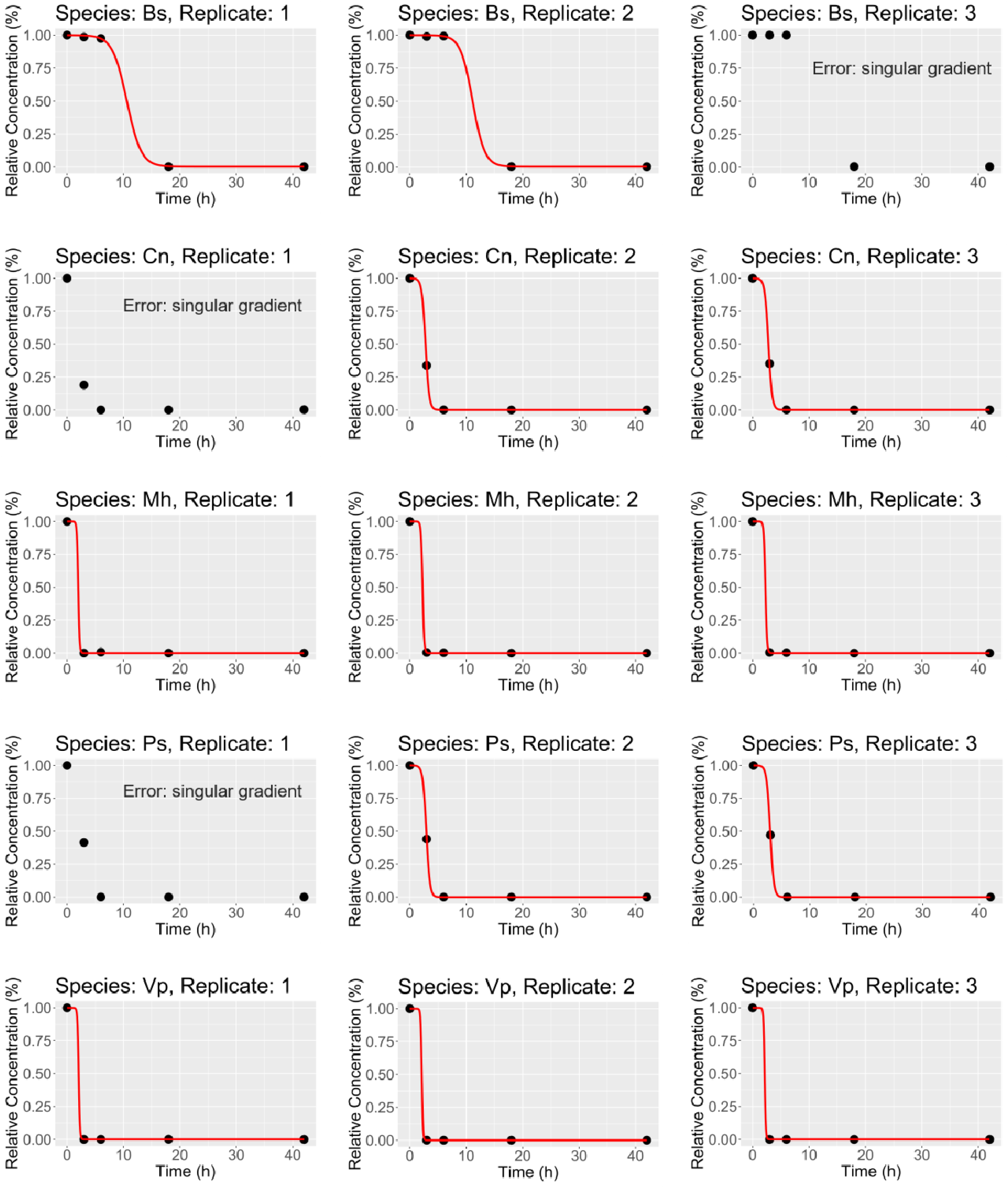
The best-fit models (red lines) and experimental data (black markers) of glutamine depletion over time using Behrends model for each of the three replicates of five bacterial monocultures. Bs*-Brevundimonas sp*., Cn*-C. necator*, Mh*-M. hominis*, Ps*-P. soli*, Vp*-V. paradoxus*. For *Brevundimonas sp*., *C. necator*, and *P. soli*, one replicate yielded singular gradient error during curve fitting, likely due to insufficient unique data points so there may be more than one optimal solution for parameter values.

**SI Fig 5.**
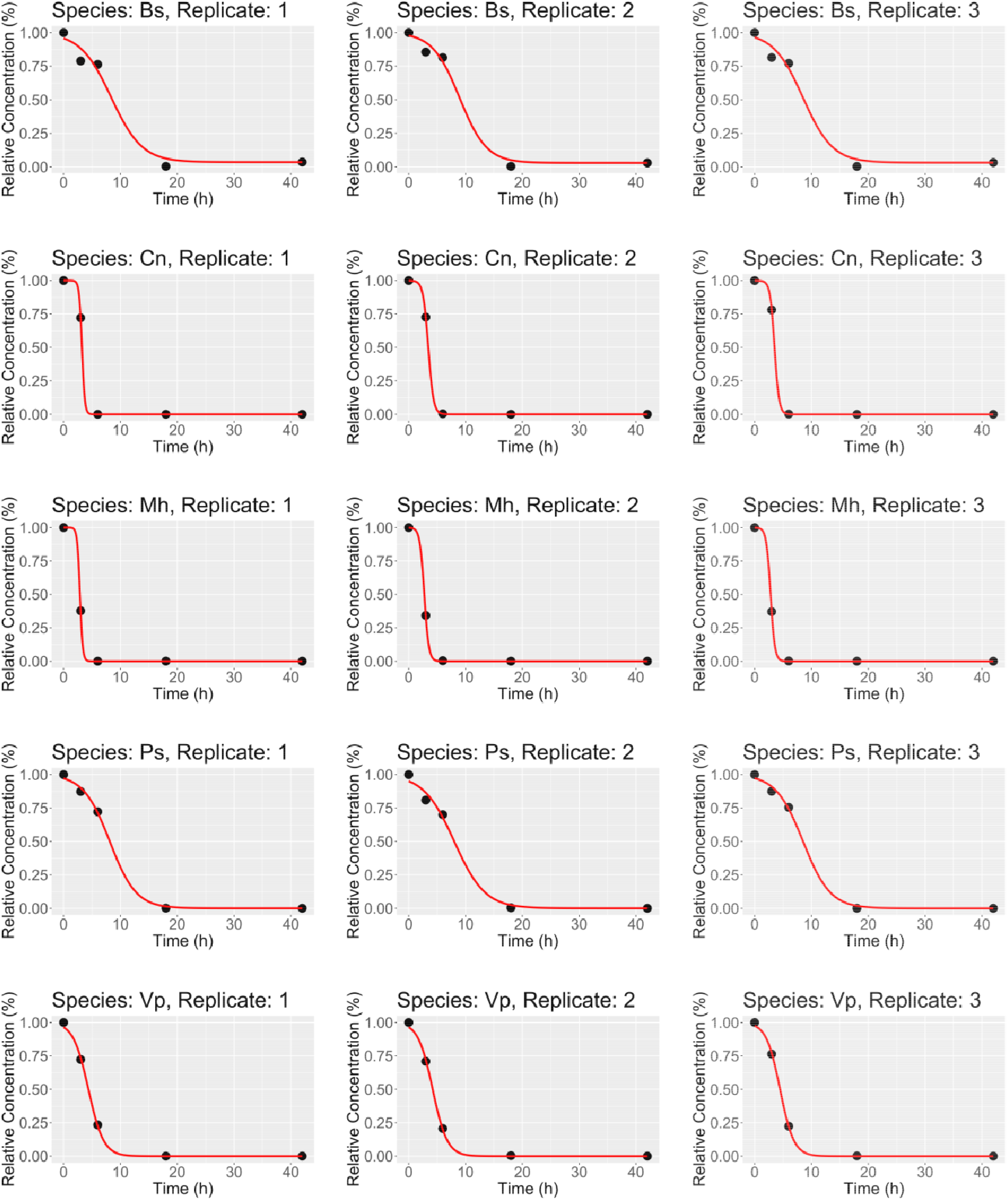
The best-fit models (red lines) and experimental data (black markers) of phenylalanine depletion over time using Behrends model for each of the three replicates of five bacterial monocultures. Bs*-Brevundimonas sp*., Cn*-C. necator*, Mh*-M. hominis*, Ps*-P. soli*, Vp*-V. paradoxus*.

**SI Fig 6.**
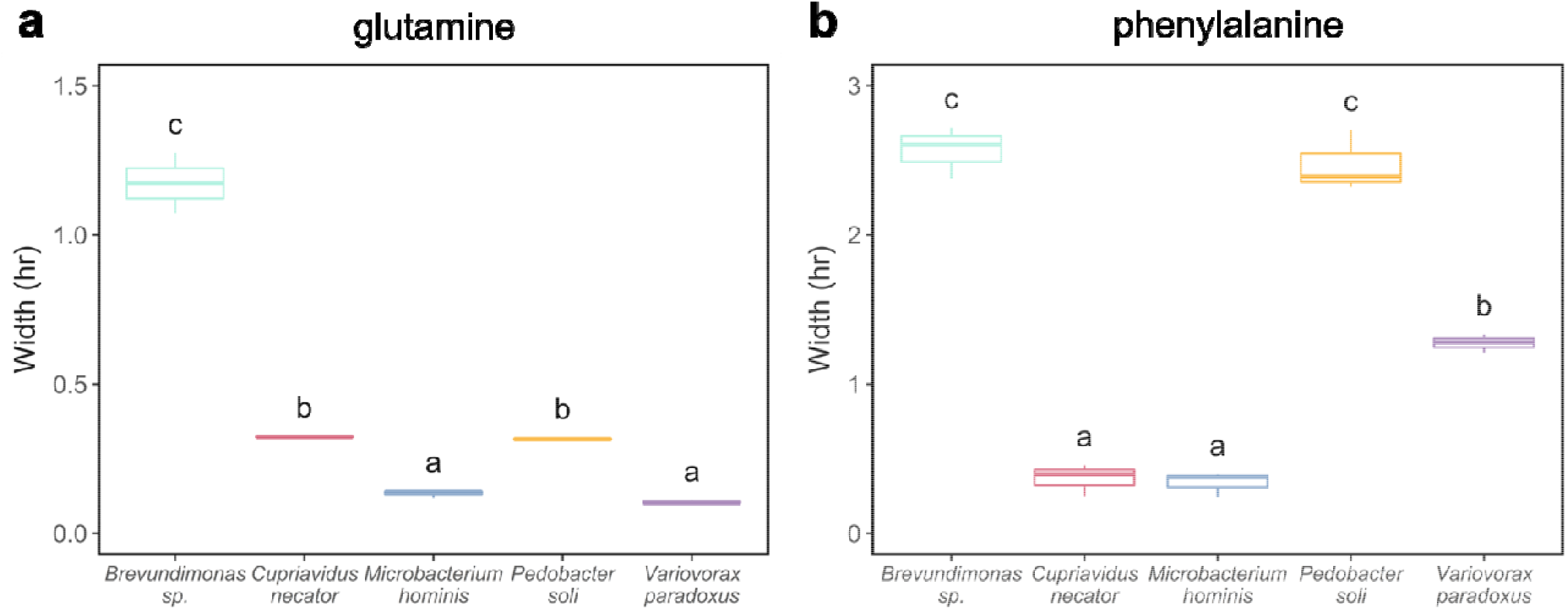
Estimated width values for use of the substrates **a**) glutamine and **b**) phenylalanine for each monoculture. Letters signify significant differences (*p* < 0.05) between species for the same substrate, using one-way ANOVA with post-hoc Tukey HSD test (n = 3, except for glutamine n = 2 for *Brevundimonas sp*., *C. necator*, and *P. soli*; SI Table 1).

**SI Fig 7.**
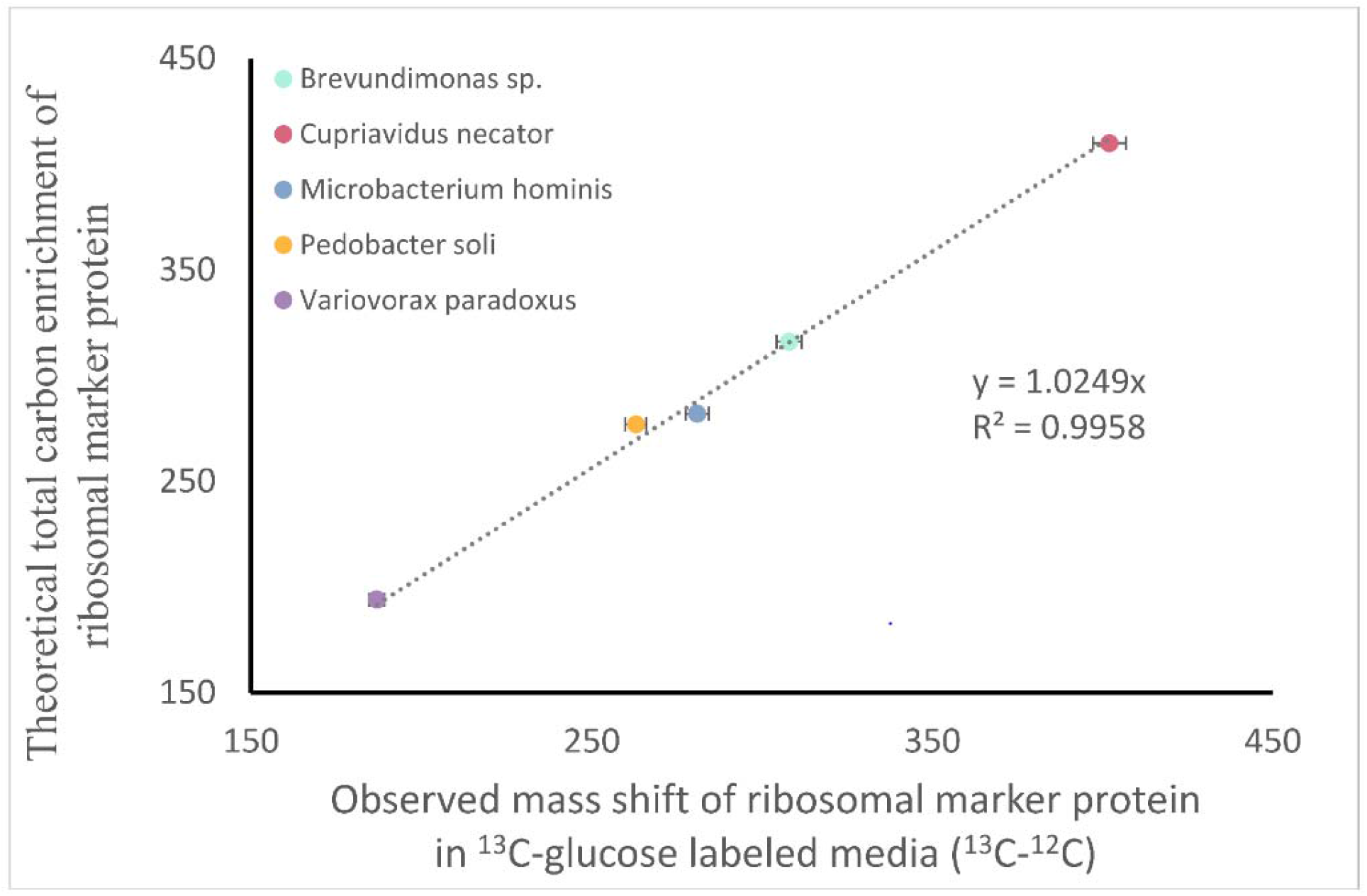
^13^C-glucose labeling study for determination of the maximal ribosomal marker protein labeling factor in monocultures (i.e., 100% enrichment). Strains were grown in ^13^C-glucose labeled media for 18 hrs, prior to MALDI-TOF MS analysis of the biological replicates (n = 4), to ensure incorporation of the substrate could be accurately tracked into the marker protein. As glucose is a preferred bacterial carbon source, this is used to provide the upper bound (100% incorporation) of ^13^C incorporation. Parallel growth experiments in unlabeled media were used to determine the lower scaling boundary (0% incorporation) for each given ribosomal marker protein. An R-squared value of 0.994 confirmed a high level of correlation between the observed mass shift, obtained from ribosomal marker protein profiling, and theoretical total carbon enrichment, calculated from the number of carbons in each protein and the natural abundance of ^13^C.

